# Transient Probability Distributions of Gene Regulatory Networks with Slow Promoter Kinetics

**DOI:** 10.1101/514547

**Authors:** M. Ali Al-Radhawi

## 1. Introduction

In a previous work [1], we have shown that, under a suitable time-scale separation hypothesis, the stationary probability mass functions of gene regulatory networks can be written as mixtures of Poisson distribution.

In this work, we study the time-evolution of the probability mass function and show that it can be approximated via similar techniques.

First, we review the relevant material from [1].

### 1.1 The Reaction Network Structure

In this paper, a GRN is formally defined as a set of nodes (genes) that are connected with each other through regulatory interactions via the proteins that the genes express. The regulatory proteins are called *transcription factors*(TFs). A TF regulates the expression of a gene by reversibly binding to the gene’s promoter and by either enhancing expression or repressing it.

The formalism we employ in order to describe GRNs at the elementary level is that of Chemical Reaction Networks (CRNs) [2]. A CRN consists of *species* and *reactions*, which we describe below.

#### Species

The species in our context consist of promoter configurations for the various genes participating in the network, together with the respective TFs expressed from these genes and some of their multimers. A configuration of a promoter is characterized by the possible locations and number of TFs bound to the promoter at a given time. If a promoter is expressed constitutively, then there are two configurations specifying the expression activity state, active or inactive. A multimer is a compound consisting of a protein binding to itself several times. For instance, dimers and trimers are 2-mers and 3-mers, respectively. If a protein forms an *n*^th^-order multimer then we say that it has a cooperativity index of *n*. If species is denoted by X, then its copy number is denoted by *X*. The set of all species is *𝒮*.

For simplicity we assume the following:

(**A1**) Each promoter can have up to two TFs binding to it.;

(**A2**) Each TF is a single protein that has a fixed cooperativity index, i.e, it cannot act as a TF with two different cooperativity indices;

(**A3**) Each gene is present with only a *single copy*.

All the above assumptions can be relaxed.

Consider the *i*^th^ promoter. The expression rate of a gene is dependent on the current configuration of its promoter. We call the set of all possible such configurations the *bindingsite set B_i_*. Each member of *B*_*i*_ corresponds to a configuration that translates into a specific species 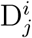 *j* ∊ *B*_*i*_. If a promoter has just one or no regulatory binding sites, then we let *B*_*i*_ = {0, 1}. Hence, the promoter configuration can be represented by *two species*: the unbound species 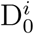 and the bound species 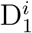. If the promoter has no binding sites then the promotor configuration species are interpreted as the inactive and active configurations, respectively. On the other hand, if the promoter has *two binding* sites then *B*_*i*_ = {00, 01, 10, 11}^1^. The first digit in a member of *B*_*i*_ specifies whether the first binding site is occupied, and the second digit specifies the occupancy of the second binding site. Hence, the promoter configuration can be represented by four species 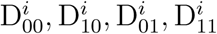. Note that in general we need to define 2^*κ*^ species for a promoter with *κ* binding sites.

The species that denotes the protein produced by the *i*^th^ gene is X_*i*_. A protein’s multimer is denoted by X_*ic*_. If protein X_*i*_ does not form a multimer then X_*ic*_:= X_*i*_.

#### Reactions

In our context, the reactions consist of TFs binding and unbinding with promoters and the respective protein expression (with transcription and translation combined in one step), decay, and *n*-merization. For each gene, we define a *gene expression block*. Each block consists of a set of *gene reactions* and a set of *protein reactions* as shown in Figure 1.

If the promoter is constitutive, i.e. it switches between two configurations autonomously without an explicitly modeled TF-promoter binding, then *B*_*i*_ = {0, 1} and the gene reactions block consists of:

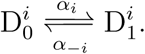

**Figure 1:**
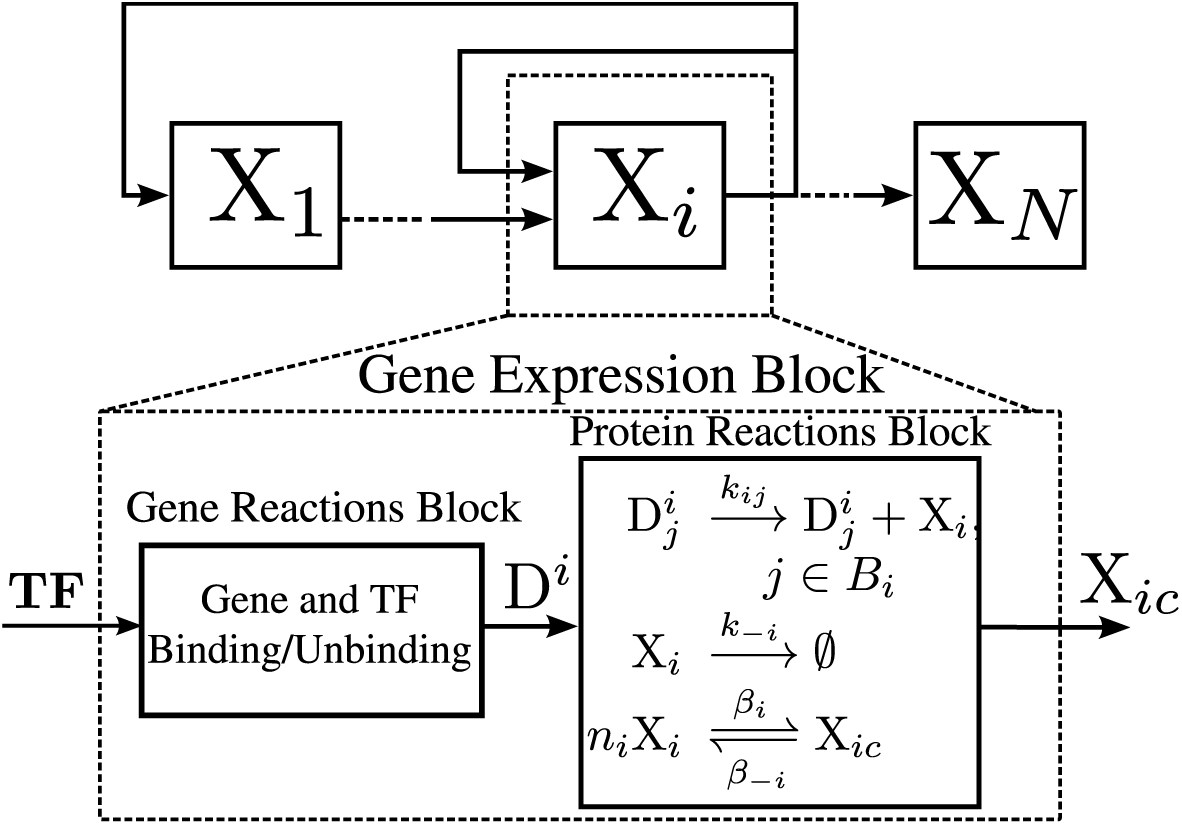
A Gene Expression Block. A GRN that consists of gene expression blocks. A block consists of a gene reactions block and a protein reactions block. The gene reactions are described in the text. **TF** is a vector of TFs which can be monomers, dimers, or higher order multimers. D^*i*^ is a vector whose components consist of the 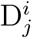’s. The dimension of **TF** is equal to the number of binding sites of the gene.

We refer to 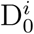 and 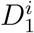 as the *inactive* and *active* configurations, respectively. If the promoter has one binding site, then also *B*_*i*_ = {0, 1} and the gene reactions block consists of just two reactions:

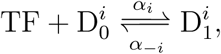

where 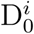 and 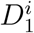 are are the promoter configurations when *unbound* and *bound* to the TF, espectively. Note that we did not designate a specific species as the active one since it depends on whether the TF is an activator or a repressor. Specifically, when TF is an activator, 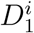 will be the active configuration and 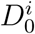 will be the inactive configuration, and vice versa when TF is a repressor. Finally, if the promoter has two TFs binding to it, then they can bind *independently, competitively*, or *cooperatively*. Cooperative binding is discussed in SI-*§*4.3.1. If they bind independently, then the promoter has two binding sites. Hence, *B*_*i*_ = {00, 01, 10, 11} and the gene block contains the following *gene reactions*:

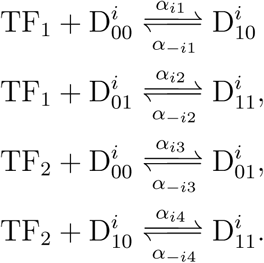

The activity of each configuration species is dependent on whether the TFs are activators or repressors, and on how they behave jointly. This can be characterized fully by assigning a production rate for each configuration as will be explained below. In the case of competitive binding, two different TFs compete to bind to the same location. This can be modeled similarly to the previous case except that the transitions to 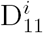, i.e. the configuration where both TFs are bound, are not allowed. Hence, the gene reactions block will have only the first and third, and the binding set reduces to *B*_*i*_ = {00, 01, 10}.

We assume that RNA polymerase and ribosomes are available in high copy numbers, and that we can lump transcription and translation into one simplified “production” reaction. The rate of production is dependent on the promoter’s configuration. So for each configuration 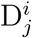, *j* ∊ *B*_*i*_ the *production* reaction is:

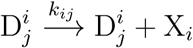

where the kinetic constant *k*_*ij*_ is a non-negative number. The case *k*_*ij*_ = 0 means that when the promoter configuration is 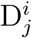 there is no protein production, and hence 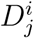 is an inactive configuration. The promoter configuration can be ranked from the most active to the least active by ranking the corresponding production kinetic rate constants.

Consequently, the character of a TF is manifested as follows: if the maximal protein production occurs at a configuration with the TF being bound we say that the TF is *activating*, and if the reverse holds it is *repressing*. And, if the production is maximal with multiple configurations such that the TF is bound in some of them and unbound in others then the TF is neither repressing nor activating.

We model decay and/or dilution as a single reaction:

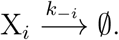

The expressed proteins can act as TFs. They may combine to form dimers or higher order multimers before acting as TFs. The numbers of copies of the TF needed to form a multi-mer is called the *the cooperativity index* and we denote it by *n*. Hence, we model the cooperativity reactions as given in Figure 1 as follows (called the *n-imerization* reactions):

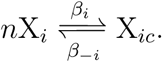

If the cooperativity index of X_*i*_ is 1, then the species X_*ic*_:= X_*i*_, and the multimerization reaction becomes empty.

### 1.2 The Chemical Master Equation

A generic reaction R_*j*_ *∈ℛ* takes the form:

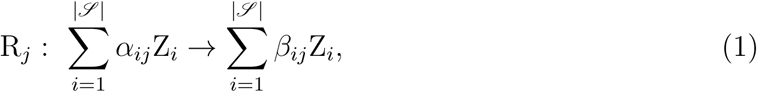

where *Z_i_ ∈𝒮, α_ij_*, and *β*_*ij*_ are positive integers. The reactions that we consider are limited to at most two reactants. The reverse reaction of R_*j*_ is the reaction in which the products and reactants are interchanged. If the network contains both the reaction and its reverse then we use the short-hand notation to denote both of them as

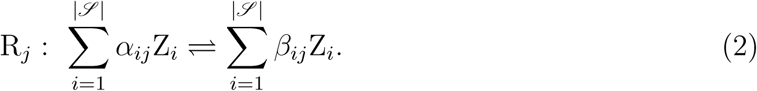

The stoichiometry of a CRN can be summarized by a *stoichiometry matrix* Γ which is defined element-wise as follows:

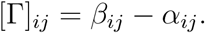

The columns of the stoichiometry matrix *γ*_1_, .., *γ|ℛ|* are known as the stoichiometry vectors. We say that a nonzero nonnegative vector *d* gives a *conservation law* for the stoichiometry if *d*^*T*^ Γ = 0.

#### Kinetics

The kinetics of the network quantify the speed of transformation from reactants into products whenever a reaction occurs. In order to keep track of molecule counts, each species Z_*i*_ *∈𝒮* is associated with a copy number *z_i_ ∈*𝕫_*≥*0_.

To each reaction R_*j*_ one associates a propensity function *R*_*j*_. Assuming a homogeneous well-stirred isothermal medium with a fixed volume, the most common model of propensities, which we use, is the *Mass-Action Kinetics* which is derived from the principle that the likelihood of two reactant molecules colliding and reacting is proportional to their copy numbers. If R_*j*_ has a single reactant species *Z*_*i*_ with stoichiometry coefficient *α*_*ij*_, then [3]:

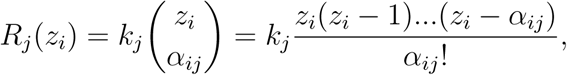

where *k*_*j*_ is a kinetic rate constant. Note if *α*_*ij*_ = 1, then *R*_*j*_(*z*_*i*_) = *k_j_z_i_*.

#### Dynamics

The dynamics of the network refers to the manner in which the *state* evolves in time, where the state 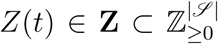 is the vector of copy numbers of the species of the network at time *t*. Since the collision of molecules is random in nature, the time-evolution of states is described mathematically by a stochastic process. The standard stochastic model for a CRN is that of a continuous Markov chain. Let **Z** denote the state space. Consider a time *t* and let the state be *Z*(*t*) = *z ∈* **Z**. Then, the probability that the *j*^th^ reaction fires in an interval [*t, t* + *δ*] is *R*_*j*_(*z*)*δ* + *o*(*δ*). If R_*j*_ fires, then the states changes from *z* to *z* + *γ*_*j*_, where *γ*_*j*_ is the corresponding stoichiometric vector.

As *Z* is a stochastic process we are interested in characterizing its qualitative behavior given by the joint probability distribution *p*_*z*_(*t*) = Pr[*Z*(*t*) = *z|Z*(0) = *z*_0_] for any given initial condition *z*_0_. The time-evolution of the probability distribution can be shown [3] to be given by a system of linear ordinary differential equations known as the *forward Kolmogorov equation* or the *Chemical Master Equation*, given by:

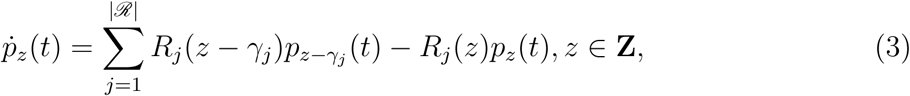

where *γ*_1_, ..*, γ_|ℛ|_* are the columns of the stoichiometry matrix.

Since our species are either gene species or protein species, we split the stochastic process *Z*(*t*) into two subprocesses: *the gene process D*(*t*) and *the protein process X*(*t*), as explained below.

Consider the *i*^th^ gene. For each configuration species 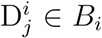, let 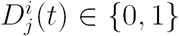 denote its occupancy, i.e. If 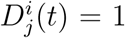, then at time *t* the *i*^th^ gene is in a configuration *j ∈ B_i_*. It can be seen from gene reactions that the network always has a conservation law supported on 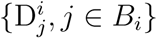, so that:

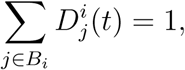

which reflects the physical constraint that the promoter can be in only one configuration at any given time.

This conservation law enables us to introduce an equivalent reduced representation. For each gene we define one process *D*_*i*_ such that *D*_*i*_(*t*) *∈ B_i_*. *D*_*i*_(*t*) = *j* if and only if 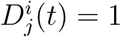. Collecting these into a vector, define the gene process *D*(*t*):= [*D*_1_(*t*), *…, D_N_* (*t*)]^*T*^ where 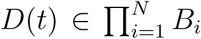. The *i*^th^ gene can be represented by *|B_i_|* states, so 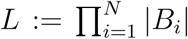 is the total number of promoter configurations in the GRN. With abuse of notation, we write also *D*(*t*) *∈* {0, ..*, L -* 1} in the sense of the bijection between {0, ..*, L -* 1} and 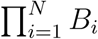 defined by interpreting *D*_1_*…D_N_* as a binary representation of an integer. Hence, *d ∈* {0, ..*, L -* 1} corresponds to (*d*_1_, *…, d_N_*) *∈ B*_1_ *× .. × B_N_* and we write *d* = (*d*_1_, ..*, d_N_*).

Since each gene expresses a corresponding protein, we define *X*_*i*1_(*t*) *∈*𝕫_*≥*0_, i = 1, ..*, N* protein processes. If the multimerized version of the *i*^th^ protein participates in the network as an activator or repressor then we define *X*_*ic*_(*t*) as the corresponding multimerized protein process, and we denote *X*_*i*_(*t*):= [*X*_*i*1_(*t*), *X*_*ic*_(*t*)]^*T*^. If there is no multimerization reaction then we define *X*_*i*_(*t*):= *X*_*i*1_(*t*). Since not all proteins are necessarily multimerized, the total number of protein processes is *N ≤ M ≤* 2*N*. Hence, the *protein process* is 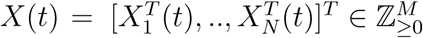 and the state space can be written as 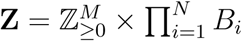.

Consider the joint probability distribution:

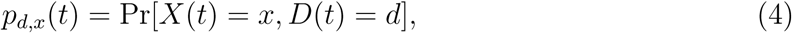

which represents the probability at time *t* that the protein process *X* takes the value 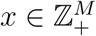 and the gene process *D* takes the value *d ∈*{0, ..*, L -*1}. Recall that *x* is a vector of copy numbers for the protein processes while *d* encodes the configuration of each promoter in the network. Then, we can define for each fixed *d*:

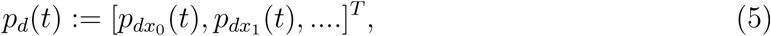

representing the vector enumerating the probabilities (4) for all values of *x* and for a fixed *d*, where *x*_0_, *x*_1_, .. is an indexing of 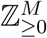. Note that *p*_*d*_(*t*) can be thought of as an infinite vector with respect to the aforementioned indexing. Finally, let

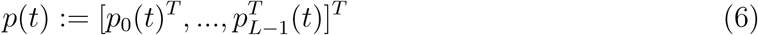

representing a concatenation of the vectors (5) for *d* = 0, ..*, L -*1. Note that *p*(*t*) is a finite concatenation of infinite vectors.

#### A gene regulatory network

Consider a set of *N* genes, binding sets 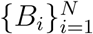, and kinetic constants *k*_*j*_’s. A *gene expression block*, as shown in Figure 1, is a set of gene reactions and protein reactions as defined above. Each gene block has an output that is either the protein or its *n*-mer, and it is designated by X_*ic*_. The input to each gene expression block is a subset of the set of the outputs of all blocks. Then, a GRN is an arbitrary interconnection of gene expression blocks (Figure 1). SI*§*4 defines a more general class of network that we can study. A directed graph can be associated with a GRN as follows. Each vertex corresponds to a gene expression block. There is a directed edge from vertex A to vertex B if the output of A is an input to B. For simplicity, we assume the following: (**A4**) The graph of gene expression blocks is connected. Note that if A4 is violated, our analysis can be applied to each connected component.

#### Time-Scale Separation

We assume that the gene reactions are considerably slower than the protein reactions. In order to model this assumption, we write the kinetic rates of gene reactions in the form *εk_j_*, where 0 *< ε «*1 and assume that all other kinetic rates (for protein production, decay and multi-merization) are *ε*^-1^-times faster.

### 1.3 Decomposition of the CME

It is crucial to our analysis to represent the linear system of differential equations given by the CME as an interconnection of weakly coupled linear systems. To this end, we present the appropriate notation in this subsection.

Consider the joint PMF: *p_d,x_*(*t*) = Pr[*X*(*t*) = *x, D*(*t*) = *d*], which represents the probability at time *t* that the protein process *X* takes the value 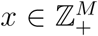 and the gene process *D* takes the value *d ∈*{0, ..*, L-*1}. Recall that *x* is a vector of copy numbers for the protein processes while *d* encodes the configuration of each promoter in the network. Then, we can define for each fixed 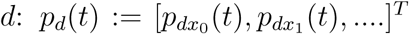, representing the vector enumerating the joint probabilities for all values of *x* and for a fixed *d*, where *x*_0_, *x*_1_, .. is an indexing of 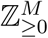. Note that *p*_*d*_(*t*) can be thought of as an infinite vector with respect to the aforementioned indexing. Finally, let

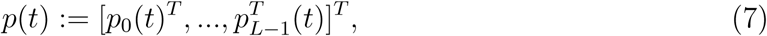

representing a concatenation of the vectors *p_d,x_*(*t*) for *d* = 0, ..*, L -*1. Note that *p*(*t*) is a finite concatenation of infinite vectors. The joint stationary PMF 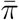 is defined as the following limit, which we assume to exist and is independent of the initial PMF: 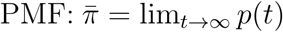. Note that 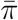 depends on *ε*.

Consider a given GRN. The CME is defined over a countable state space **Z**. Hence, the CME can be interpreted as an infinite system of differential equations with an infinite infinitesimal generator matrix Λ which contains the reaction rates.

Consider partitioning the PMF vector as in (7). Recall that reactions have been divided into two sets: slow gene reactions and fast protein reactions. This allows us to write Λ as a sum of a slow matrix 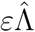 and a fast matrix 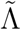, which we call a fast-slow decomposition. Furthermore, 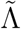 can be written as a block diagonal matrix with *L* diagonal blocks which correspond to conditioning the MC on a specific gene state *d*. This is stated in the following basic proposition (see [1] for the proof):

#### Proposition 1.

Given a GRN. Its CME can be written as

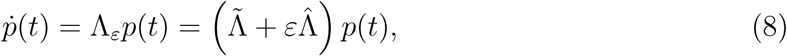

where 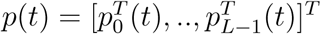, and

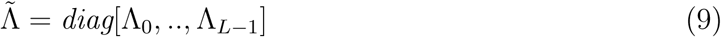

where 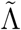 is the fast matrix, 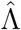is the slow matrix, and Λ_0_, .., Λ_L-1_ are stochastic matrices.

### 1.4 Conditional MCs

For each *d*, consider modifying the MC *Z*(*t*) defined in the previous section by replacing the stochastic process *D*(*t*) by a deterministic constant process *D*(*t*) = *d*. This means that the resulting MC does not describe the gene process dynamics, it only describes the protein process dynamics *conditioned on d*. Henceforth, we refer to the resulting MC as the *MC conditioned on d.* The infinitesimal generator of a MC conditioned on *d* is denoted by Λ_*d*_, and is identical to the corresponding block on the diagonal of 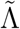 as given in (9). In other words, fixing *D*(*t*) = *d ∈*{0, ..*, L-*1}, the dynamics of the network can be described by a CME:

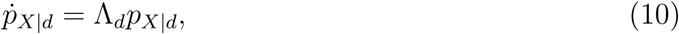

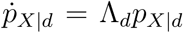, where *p_X|d_* is a vector that enumerates the conditional probabilities *p_x|d_* = Pr[*X*(*t*) = *x | D*(*t*) = *d*] for a given *d*. The conditional stationary PMF is denoted by: 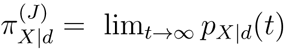, where (*J*) refers to the fact that it is joint in the protein and multimerized protein processes. Note that 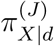 is independent of *ε*. This notion of a conditional MC is useful since, at the SPK limit, *D*(*t*) stays constant. It can be noted from (9) that when *ε* = 0 the dynamics of *p*_*d*_ decouples and becomes independent of 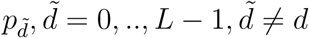.

We show below that each conditional MC has a simple structure. Fixing the promoter configuration *D*(*t*) = *d* = (*d*_1_, ..*, d_N_*), the network consists of *uncoupled* birth-death processes. So for each *d*_*i*_, the protein reactions of production and dimerization corresponding to the *i*^th^ promoter can be written as follows without multimerization: 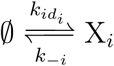, where the subscript *id*_*i*_ refers to the production kinetic constant corresponding to the configuration species 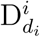 or, if there is a multimerization reaction, it takes the form: 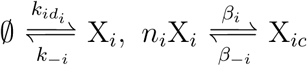. Note that the stochastic processes X_*i*_(*t*), *i* = 1, ..*, N* conditioned on *D*(*t*) = *d* are independent of each other. Hence, the conditional stationary PMF 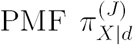 can be written as a product of stationary PMFs and the individual stationary PMFs have Poisson expressions. The following proposition gives the analytic expression of the conditional stationary PMFs: (see [1] for proof)

#### Proposition 2.

Fix d ∈ {0, .., L - 1}. Consider (10), then there exists a conditional stationary PMF 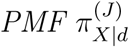 and it is given by

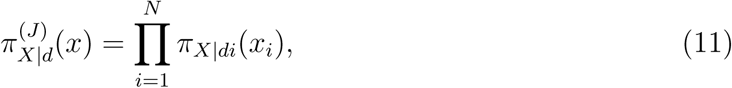

where

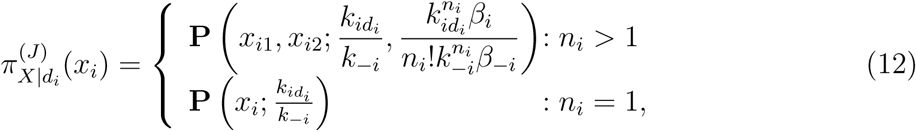

where (J) refers to the joint PMF in multimerized and non-multimerized processes, x_i1_ refers to the copy number of X_i_, while x_i2_ refers to the copy number of X_ic_, 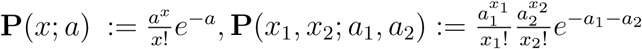.

#### Remark 1.

The conditional PMF in (11) is a joint PMF in the protein and multimerized protein processes. If we want to compute a marginal stationary PMF for the protein process only, then we average over the multimerized protein processes X_ic_, i = 1, .., N to get a joint Poisson in N variables. Hence, the formulae (11)-(12) can be replaced by:

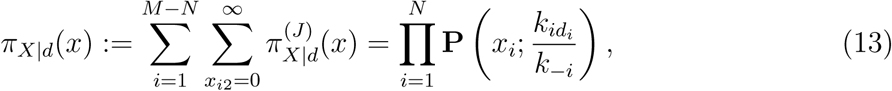

where M-N is the number of n-merized protein processes, and π_X|d_ is the marginal stationary PMF for the protein process.

Recall the conditional Markov chains with the associated infinitesimal generators as in Eq. (4). By the assumption for each *d ∈* {0, ..*, L -* 1} there exists a unique *π*_*X|d*_ such that: Λ*_d_π_X|d_* = 0, *π*_*X|d*_ *>* 0, and Σ*x π_X|d_*(*x*) = 1. Recall that *π*_*X|d*_ is the stationary distribution of the Markov chain conditioned on *D*(*t*) = *d*.

Defining the extended conditional distributions for *d* = 0, ..*, L -* 1 as:

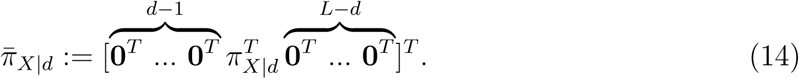

The stationary distribution above can be interpreted as a function as follows: 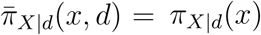, and 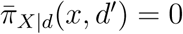 when *d ′ ≠d*.

### 1.5 Stationary Distribution at the limit of slow promoter kinetics

Recall the slow-fast decomposition of the CME in (8) and the joint stationary PMF 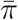. In order to emphasize the dependence on *ε* we denote 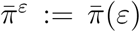 Hence, 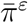 is the unique stationary PMF that satisfies 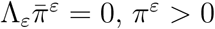, and 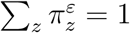, where the subscript denotes the value of the stationary PMF at *z.*

Our aim is to characterize the stationary PMF as *ε→* 0. Writing 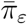 as an asymptotic expansion to first order in terms of *ε*, we have

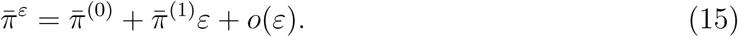

Singular perturbation techniques can be used to derive the following theorem [1]:

#### Theorem 3.

Consider a given GRN with L promoter states with the CME (8) Writing (15), then the joint stationary 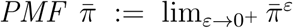 can be written as: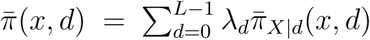, where λ = [λ_0_, .., λ_L-1_]^T^ the principal normalized eigenvector of:

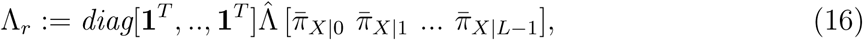

where 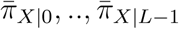 are the extended conditional stationary PMFs defined as: 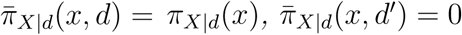 when d ′ ≠d.

## 2 Transient Dynamics of the CME

In [1], we have been interested in computing the analytical expression for the stationary PMF at the SPK limit. In this section, we use the same techniques to write the transient dynamics after an initial time interval (called *the boundary layer* in the singular perturbations literature [4].)

Recall that Proposition 1 provides a decomposition of the CME as follows:

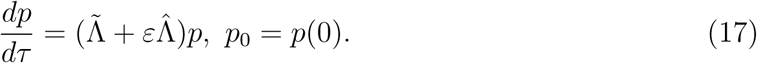

The effect of the slow gene reactions matrix 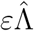 in the model above takes a relatively long time to be active. Because of this, one calls *τ* the “*fast*” time-scale. So in order to for the effect of 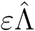 to manifest sooner, we define the “slow time-scale” *t* = *ετ*. The Master Equation can be written as follows:

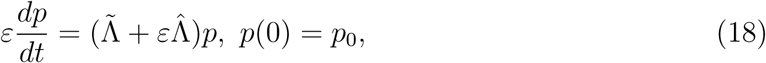

or equivalently as 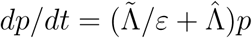.

Let *p_ε_*(*t*) be the solution of (18). Writing:

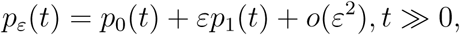

and substituting the above expansion in (18), we obtain the following equation from the zeroth-order term:

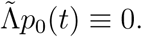

Hence, the distribution *p*_0_ will quickly approach the kernel space of the fast matrix, i.e 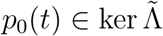, which is spanned by the extended conditional distributions 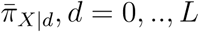 as given in (14). Hence, we obtain:

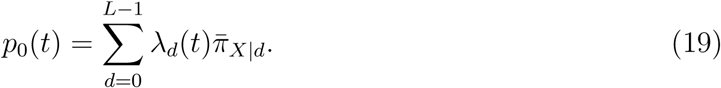

The first order term in this expression gives the following equation:

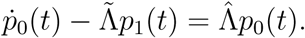

Recall that 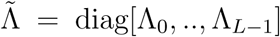, and that each Λ_*d*_ is an infinitesimal generator which satisfies **1**^*T*^ Λ_*d*_ = 0. Hence, if we pre-multiply the equation above by 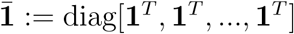 and substitute (19) we obtain:

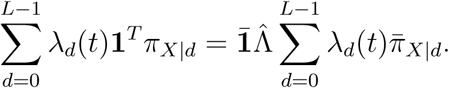

Since **1***^T^ π_X|d_* = 1 and recalling the definition of the reduced infinitesimal generator Λ_*r*_ (16), we get the following finite dimensional linear system:

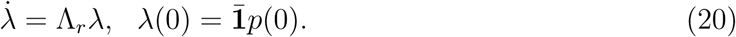

This is an autonomous linear system and the solution can be written as a matrix exponential as follows: 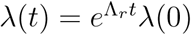.

Hence, as *ε→*0 the solution of the master equation is approximated by the following expression after the boundary layer:

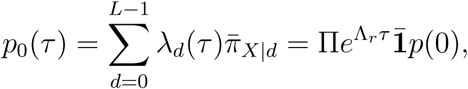

where 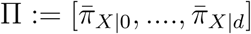.

In the slow time-scale, we can write

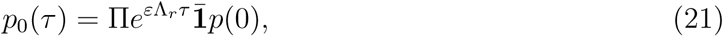

which is consistent with the analysis carried out in [5] for the finite projection of the CME.

In order to estimate the initial boundary layer, let 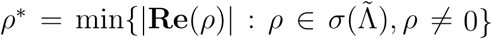, where *σ*(.) is the spectrum operator. Assuming that *ε < ρ^*^*, it can be seen that the approximation *p*(*τ*) = *p*_0_(*τ*)+ *o*(*ε*) is valid for *τ > T* where *T* is proportional to - ln *ε/ρ^*^*. In order to compare to the main results of [1], recall that Λ_*r*_ itself is an infinitesimal generator, hence 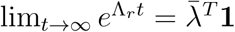, where 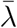 is the principal eigenvector of Λ_*r*_. Hence,

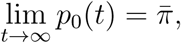

where 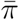 is the stationary PMF in the statement of Theorem 3. Hence, a generalization of

Corollary 4 in [1], can be stated as:

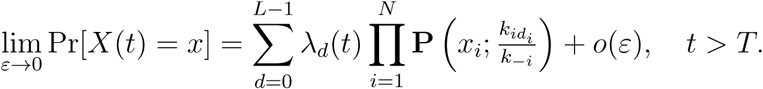

## Example: The Self-Repressing Gene

We now revisit the self-repressing gene:

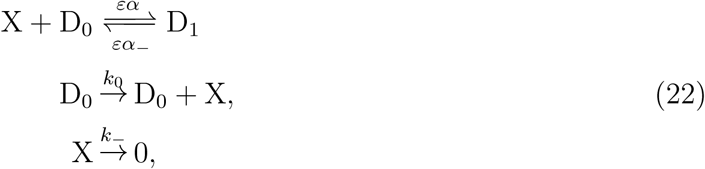

which was studied in Figure 1 in [1].

Using (16), the reduced generator can be written as:

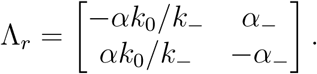

The stationary PMF is as follows:

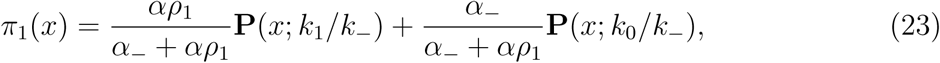

Where

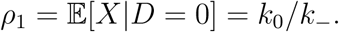

We study the case *ε* = 0.1. Figure 2-a shows how the stationary distribution evolves in time. It can be seen that the distribution becomes bimodal at the two predicted modes at around *t* = - (ln *ε*)*/ρ^*^≈* 2.35, so it approaches the the kernel of 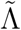 quickly. Then, the slow dynamics given by (20) drive the time evolution of the weights of the two modes to their steady state values. Figure 2-b depicts evolution of the probability of Pr[*X*(*t*) = 40], and compares the exact and approximated solutions. It can been seen that the two solutions match after an initial boundary layer.

**Figure 2:**
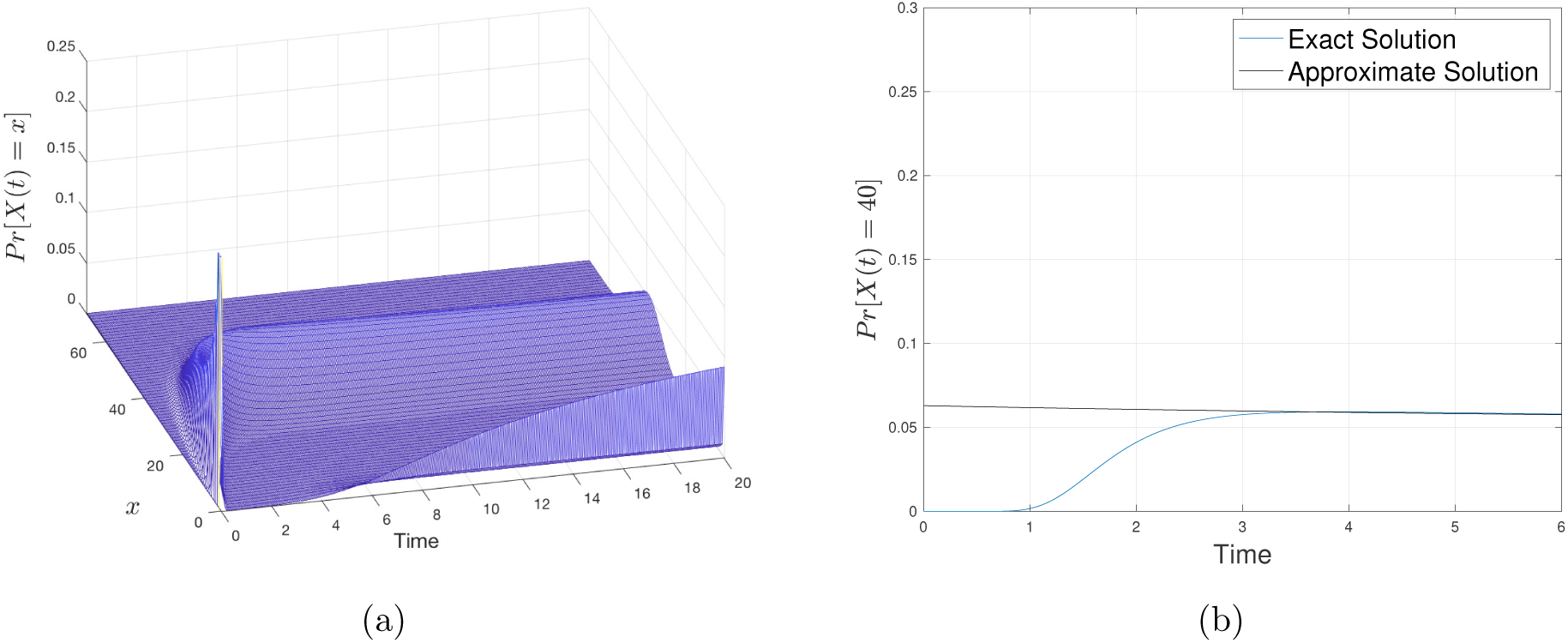
Time evolution of the PMF for a self-repressing gene with. *ε* = 0.1. (a) The full-state time evolution of the probability distribution for self-repressing gene with the initial condition Pr[*X*(0) = 0, *D*(0) = 0] = 1. (b) The time evolution of *Pr*[*X*(*t*) = 40] compared between the finite state projection solution [5], and the singular perturbation approximation (21). The full trajectories have been generated by a finite projection numerical solution of the CME. Both figures use the network with parameters *α* = *ε/*200, *α*_-_ = *ε, k*_0_ = 40, *k*_-_ = 1.

## Footnotes

1 We interpret the elements of the binding set as integers in binary representation.

